# Robust and Adaptive Deep Model Ensemble Framework Fine-tuned by Structural Information for Drug-Target Interactions

**DOI:** 10.1101/2023.10.20.563031

**Authors:** Jinhang Wei, Linlin Zhuo, Xiangzheng Fu, Junmin Zhang, Xiangxiang Zeng, Quan Zou

## Abstract

In the fields of new drug development and drug repositioning, drug-target interactions (DTI) play a pivotal role. Although deep learning models have already made significant contributions in this domain, the state-of-the-art models still exhibit shortcomings in predictive performance and issues of false-negative errors. Based on these observations, we constructed a streamlined yet effective base learner model. With our designed adaptive feature weight network, the model can capture key features within drugs (targets). Furthermore, by cross-partitioning the training data, multiple base learners are integrated into a powerful ensemble model named EADTN. The performance of the model is further enhanced as the number of base learners increases. Additionally, we employed a single-linkage clustering algorithm to cluster drugs and proteins and leveraged this clustering information to fine-tune the base learners, which elevates the value of EADTN in real-world applications like drug repositioning and targeted drug development. Our designed substructure importance ranking method also demonstrates the model’s exceptional capability to recognize key substructures. Benefiting from the model’s low generalization error capability, we successfully identified false-negative samples within the dataset, revealing new interaction relationships. Experimental results indicate that EADTN consistently outperforms existing state-of-the-art models across multiple datasets. More importantly, the ensemble learning and clustering fine-tuning approaches adopted by our model offer a fresh perspective for related fields.

## 1 Introduction

Drug-target interaction (DTI) is a pivotal issue in drug development, where the efficacy of a drug often depends on its interaction with biomolecules such as proteins and DNA [1]. Therefore, accurately predicting the interactions between drugs and their target sites holds significant scientific and clinical value [2]. However, experimentally measuring DTIs demands considerable time and resource investments, hindering its application in large-scale screening and drug repurposing. As a result, establishing effective computational methods to predict interactions between drugs and target proteins has become a major challenge in drug research [3].

In recent years, with advancements in scientific research, there’s a deeper understanding of drug-target interactions. For instance, the application of molecular docking and virtual screening techniques allows us to simulate the interactions between drugs and target proteins [4]. Through these methods, researchers can screen potential drug candidates from millions or even billions of compounds and predict their binding modes and affinities with the target proteins. This not only greatly boosts the efficiency of drug development but also provides profound insights into the functions of target proteins and the mechanisms of drug action [5]. For example, Beck et al. [6] employed computational methods to sift through extensive datasets, predicting that Azanavir, Remdesivir, and Kaletra could inhibit SARS-CoV-2. Mahmoud and colleagues [7] utilized computational molecular modeling to screen over 2000 FDA-approved drugs with established safety profiles and found that Atovaquone, Methylene Blue, and Valinomycin exhibited significant antiviral activities against SARS-CoV-2. Xia et al. [8] conducted a case study on Melphalan (a key drug in anti-tumor therapy) using deep learning, revealing that Melphalan might activate HMOX1, inhibit CDC20, and interact with MYC, consistent with previous research findings. These instances underline the importance and effectiveness of computational methods in drug development. Therefore, the creation of precise and efficient computational approaches can significantly narrow down the scope of wet-lab experiments, reduce Research and Development costs, and propel advancements in drug research.

Within the realm, significant advancements have been made in DTI prediction using machine learning and deep learning technologies. These predictive models forecast potential interactions by learning molecular features of drugs and their targets. For instance, DrugBAN [9] employs an interpretable bilinear attention network based on drug molecular graphs and target protein sequences, capturing the interplay between drugs and their targets. MolTrans [10] relies on a substructure pattern mining algorithm and interaction modeling module, utilizing an augmented transformer encoder to predict DTI relations. GraphDTA [11] employs graph neural networks to represent drugs as molecular graphs, enhancing the model’s performance in predicting drug-target affinity. DeepConv-DTI [12] utilizes convolutional neural networks [13] (CNN) to grasp generalized protein classes’ local residue patterns, predicting drug-target interactions. Nonetheless, despite the notable advancements made by deep learning techniques in DTI prediction, various models may have different representations and understandings of drug-target interactions, potentially limiting them in addressing certain types of DTI challenges. To further enhance the accuracy and robustness of DTI prediction, employing ensemble learning could be a feasible approach.

Ensemble learning is a machine learning technique aimed at improving predictive performance by integrating multiple models [14]. The basic concept involves generating a set of base learners and aggregating their predictions through mechanisms like voting, averaging, or other combination strategies. A foundational premise of ensemble learning is that a collective effort often surpasses the performance of any individual model, as they can compensate for each other’s shortcomings, reduce overfitting, and increase robustness [14]. One critical goal of ensemble learning is to bolster predictive performance by enhancing model diversity. Diversity can be achieved either by employing diverse model types or by training the same type of model with different datasets [15]. Although ensemble learning has demonstrated its prowess across multiple machine learning tasks [16], its application remains relatively limited in DTI prediction tasks.

Given the aforementioned context and challenges, we introduce a novel model, termed EDATN. In EDATN, base learners employ convolutional neural networks to extract features from protein primary structures and drug SMILES. These features are then independently adjusted through an adaptive feature weight network. This procedure aims to capture the potential feature diversity across different targets and drugs, enabling precise drug-target interaction predictions. Furthermore, we employ a k-fold cross-validation strategy to perturb training data, training k base learners with identical structures. This step amplifies ensemble learning’s diversity. Subsequently, we implement an average pooling strategy, amalgamating the predictions from the k base learners, yielding EDATN’s final prediction. However, we also recognize that in practical applications, like drug repositioning and target prediction [17], predictions for new interactions often need to be based on data similar in structure or properties to the target drug (or target). Therefore, we conceptualized a novel prediction scenario, which we refer to as the “intra-cluster” scenario. We use a single-linkage algorithm and base our clusters on ECFP4 fingerprints and pseudo-amino acid composition, clustering drugs and proteins. We then partition each cluster into new training and test sets at a predefined ratio. To optimize the model in this scenario, we initially train each base learner with a fixed training set. Then, based on clustering information, we split the training set into k subsets, subsequently combining n of them to yield k finetuned training sets. Each base learner undergoes fine-tuning accordingly. During DTI relation prediction, the model first discerns the target’s cluster info and then boosts the credibility of the output from the related cluster’s fine-tuned learner, leading to the final prediction. Based on experimental outcomes, this unsupervised clustering-based fine-tuning strategy can further amplify the model’s predictive performance. Our contributions can be summarized as:

1. We devised a DTI prediction model based on ensemble deep learning, named EADTN, uncovering latent DTI relations.
2. By incorporating a clustering fine-tuning strategy, the practicality of EADTN in drug repositioning and target prediction is augmented.
3. Leveraging shapley values, we conceptualized a model-agnostic substructure verification method, successfully validating EADTN’s precision in key substructure identification.
4. Through case study and in the context of potential false negatives, combining model and molecular structure verification, we unearthed new drug-target interaction relations.

## 2 Materials and methods

### 2.1 Dataset

The data used in this study primarily comes from two publicly available datasets: BindingDB [18] and BioSNAP [19].

BindingDB [18] is an open-access biomedical database related to medicinal chemistry and pharmacology, focusing primarily on the collection of interaction affinities between drug target proteins and drug-like small molecules. BindingDB accumulates experimental data from research institutions worldwide, including target names, target sequences, drug names, drug structures, and pathway information. This wealth of experimental data from BindingDB is immensely valuable for our research.

The drug-target interaction network in BioSNAP [19] is a database that contains information about the genes (i.e., proteins encoded by the genes) targeted by drugs on the US market. Within this database, drug target data comprises biotechnological drugs and dietary supplements (these drugs, on average, target 5-10 unique proteins). It lists all identified targets with physiological or pharmacological effects (not just a single primary target) and extensively indicates that many drug targets are protein complexes made up of multiple subunits or protein combinations.

Before commencing the research, we cleaned and preprocessed all the data to ensure data quality and usability. Ultimately, the BindingDB dataset included 14,643 drugs, 2,623 target proteins, and 20,674 drug-target interaction relationships (DTIs). The BioSNAP dataset comprised 4,505 drugs, 2,181 target proteins, and 13,830 DTIs.

### 2.2 EADTN framework

The overall framework of the EADTN model we proposed is illustrated in Figure 1. This model mainly consists of four modules: the CNN encoder module, the adaptive feature weighting module, the MLP decoder module, and the clustering-weighted module.

**Fig. 1:**
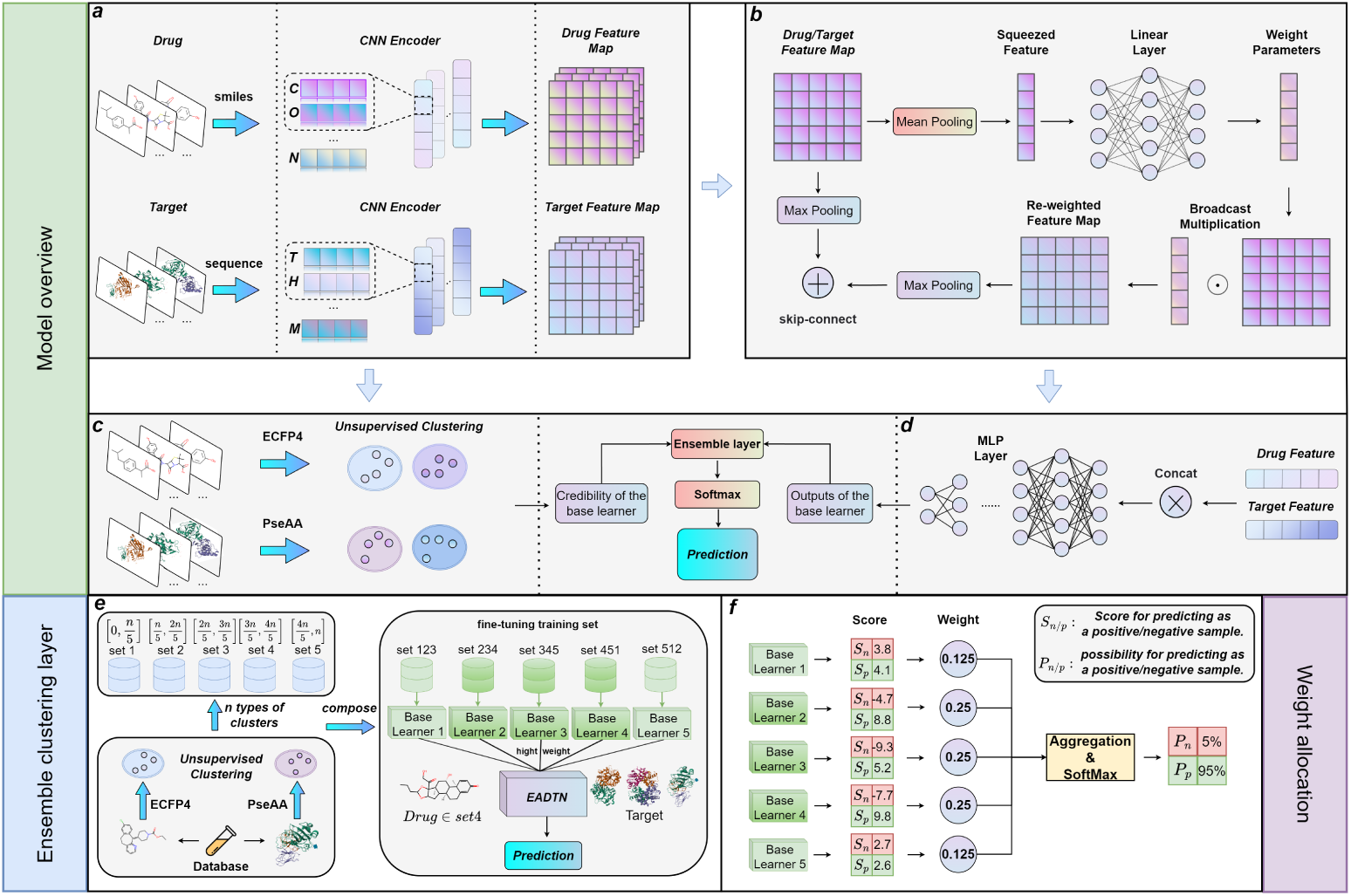
The model diagram of EADTN. a. CNN Encoder Module: Atoms and amino acids are transformed into vectors using a word embedding table. Subsequently, molecular and target characterizations are extracted using a 1D-CNN. b. Adaptive Feature Weight Module: Weights are trained based on molecular and target representations to reassess the importance of each atom and amino acid. Residual connections ensure preservation of the initial features. c. Clustering Weighting Module: Drugs and targets are subjected to similarity clustering via a single-linkage algorithm. These clustering outcomes are then harnessed to optimize the model. d. MLP Decoder Module: The molecular and target representations obtained from the adaptive weight network are decoded to forecast their potential interactions. e. Integrated Clustering Layer: Data is categorized based on clustering information, fine-tuning the base learner, and adjusting its weight during inference. f. Weight Allocation Module: This module aggregates the outputs of multiple base learners in EADTN, converting them into predictive probabilities.

The CNN encoding layer first captures the SMILES of the drug and the primary sequence of the target protein. Using a predefined embedding dictionary, characters are mapped to numerical values. Subsequently, the CNN module is employed to extract representations of the drug and the target. Ultimately, each drug (or target) is transformed into a feature map. This feature map is further forwarded to the adaptive feature weighting module.

In the adaptive feature weight module, the entire feature map’s information is compressed into a single feature vector through mean pooling first. This vector is further input into a linear layer for transformation, outputting a weight vector that will perform broadcast multiplication with the original feature map, thus forming a feature map with adjusted weights. Max-pooling and residual connections are used to integrate the information from the original feature map with that of the adjusted weight feature map, thereby forming the final feature vectors for the drug or target. Next, these obtained feature vectors for drugs and targets undergo a concatenate operation and are input into the MLP decoder module, generating a set of outputs from base learners.

As we elaborated earlier, DTI models, in practical application scenarios, often need to predict new interactions based on data that is structurally or property-wise similar to the target drug (or target). For this reason, the clustering weighting module has been specifically designed to cater to such scenarios. This module clusters drugs and proteins using the single linkage algorithm, based on the ECFP4 fingerprint and pseudo amino acid composition. This process constructs a fine-tuning training set through clustering information and pre-trains the base learners while recording which base learner belongs to each fine-tuning training set. Based on this, we can adjust the credibility of the results from multiple base learners without needing label information, only using clustering information constructed from the ECFP4 fingerprint and pseudo amino acid composition, to obtain the final results of EADTN. Therefore, our approach offers an effective way to combine the outputs of base learners with clustering weighting information, providing an efficient ensemble learning computation method for matching drugs and targets.

#### 2.2.1 CNN encoding module

In EADTN, both the drugs and the targets must be converted from SMILES and amino acid sequences to numerical vectors for the model’s intake. Initially, each character in the drug’s SMILES sequence is translated into an integer between 1 to 64. Similarly, every character in the target’s amino acid sequence is mapped to an integer between 1 to 24. To ensure consistency in sequence length, we pad zeros at the end of sequences. Subsequently, the numerical sequences of drugs and targets are fed into a trainable embedding layer, producing dense vector representations for each character. Consequently, every protein feature matrix can be denoted as *M_p_ ∈ R*^(^*^Lp∗De^*^)^ and every drug feature matrix as *M_d_ ∈ R*^(*L*^*^d ∗De^*^)^, where *L_p_* and *L_d_* represent the lengths of protein and drug sequences respectively, and *D_e_* signifies the predefined embedding size.

Following this, we employ a one-dimensional convolutional neural network (1D-CNN) to process these feature matrices. 1D-CNN exhibits robust feature extraction capabilities, especially adept at handling such sequence data by capturing local patterns and preserving spatial relationships. For both drugs and targets, the feature matrices are input into a network composed of three convolutional layers accompanied by ReLU [20] activation functions. This structure, while handling sequences, is designed to recognize multi-scale local features through its multilayered convolution. After this network processing, the feature matrix evolves into a more enriched and abstract feature representation. It’s noteworthy that this procedure does not alter the sequence length, ensuring the retention of original information while enhancing the feature description. Such attributes make 1D-CNN exceptionally well-suited for handling this type of sequence data.

### 2.3 Adaptive feature weight module

In past DTI prediction models [11, 12, 21, 22], many models did not fully consider the importance of local features. Specifically, for the acquired drug and protein feature maps, models proceeded to the next computation step merely through a simple pooling operation. However, this approach could lead to a significant loss of crucial information. Consequently, this loss of information undoubtedly impacts the model’s predictive performance and interpretability. Thus, even if the model performs well in certain scenarios, we must recognize that overlooking the importance of local features in the prediction process can undermine the model’s comprehensiveness and accuracy. As a result, we designed an adaptive feature weight module to capture the importance of local atomic features in drug molecules and local amino acid features in protein molecules. This design aims to enhance the role of essential local features, diminish the impact of less critical features, and improve the model’s performance and interpretability.

In EADTN, we first use mean pooling to obtain the global feature information of the molecule (protein), which is then fed into a linear layer without bias. Initially, we set a fixed compression ratio to determine the number of neurons in the hidden layer. Subsequently, in the output layer, we increase the feature dimension to match the input. Compressing the number of neurons in the middle layer not only reduces computational complexity when the drug (protein) sequence is too long but also prevents model overfitting while capturing global information. Expanding to the original dimension in the output layer can act as a weight for local atomic (amino acid) features, capturing more important atomic (amino acid) information from the feature map. In our work, we employed two linear layers, each utilizing the Relu activation function. The formal definition is as follows:

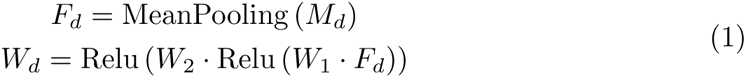

Where *M_d_* represents the drug feature matrix, *F_d_* denotes the global feature information of the drug, and Wd is the adaptive weight vector. The formula concerning the protein can be derived similarly.

Through the aforementioned formula, we successfully constructed a feature map with a reallocated weight. For the subsequent decoding process, it’s necessary to compress the feature map of the drug molecule (or protein target) into a one-dimensional feature vector. During this process, we adopted max pooling as our pooling strategy, which is consistent with our previous approach of using an adaptive weight network to adjust feature weights. The rationale behind this decision is our dedication to capturing the most influential features and describing the Drug-Target Interaction (DTI) relationship based on these dominant features. Therefore, our processing method is theoretically more persuasive, as it zeroes in on the features that have the most significant impact on the outcome, rather than the average effect of all features. While this strategy might cause certain features to be overlooked, it allows the model to focus more on critical information, thereby avoiding noise interference from excessive non-dominant features.

Next, we employ the max pooling method to compress the initial feature map and integrate it with the features reallocated by the weight network. This process essentially fuses two different sources of information. The feature vector obtained from max pooling of the initial feature map tends to preserve spatial information to a greater extent, while the feature vector processed by the weight network mainly considers relationships between substructures. Thus, this strategy not only compensates to some extent for the potential information loss caused by max pooling but also enhances the robustness of the adaptive weight network. The formulaic definition of the above operations is:

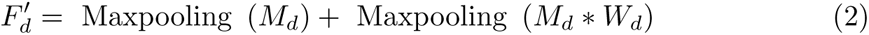

Where *∗* denotes the broadcast multiplication, *W_d_* is the weight vector outputted by the drug’s adaptive weight network, and *M_d_* is the drug feature matrix. The formula concerning the protein can be derived in a similar manner.

### 2.4 MLP decoder module

Figure 2 illustrates the complete data flow process from the adaptive feature weight module to the Multi-Layer Perceptron (MLP) decoder module. Subsequently, we concatenate the drug feature vectors with the target feature vectors. This allows us to represent both drugs and targets simultaneously within a unified vector space, thereby better capturing their intrinsic relationships and interactions.

**Fig. 2:**
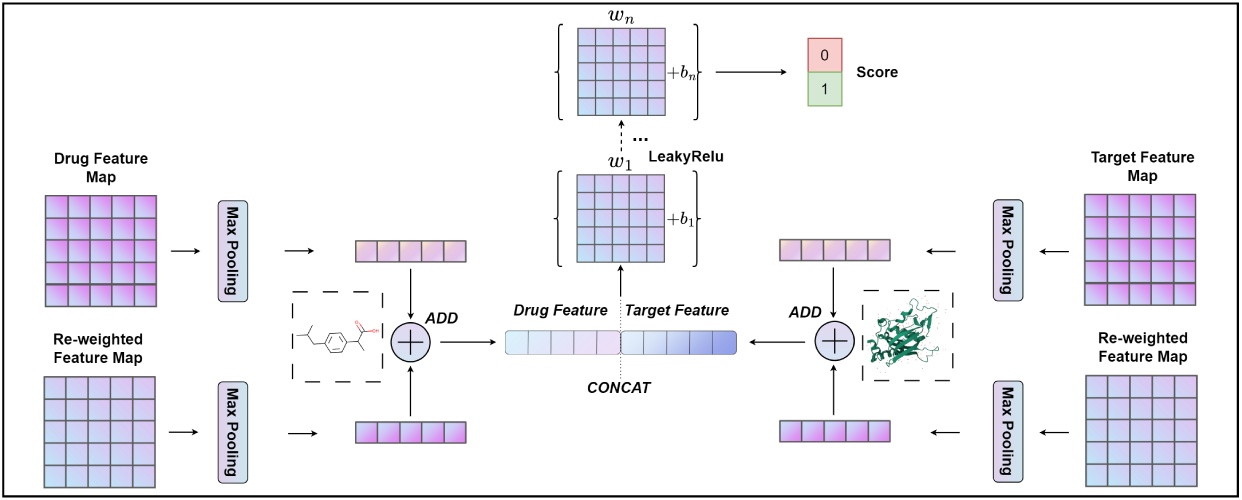
Flowchart of MLP decoder module.

Lastly, the concatenated features are fed into the MLP. After undergoing several layers of linear transformations and non-linear activation functions, a higher-level abstraction and representation of the data is achieved. To optimize the model’s generalization capability and prevent overfitting, we apply the LeakyReLU [23] activation function after each linear transformation, aiming to enhance the model’s fitting and expressiveness. Compared to the ReLU activation function, LeakyReLU retains the positive signal while also allowing the transmission of negative signals. Ultimately, the output layer produces scores of 0 (indicating no interaction) and 1 (indicating interaction). These scores are then used in the ensemble learning layer, and the base learner prediction process can be expressed as:

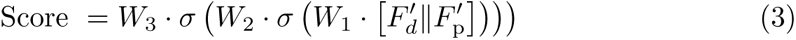

where*||* denotes the concatenate operation, and *σ* represents the LeakyReLU activation function.

### 2.5 Cluster weighting module

In this study, we independently clustered drug compounds and target proteins from the BindingDB and BioSNAP datasets, aiming to construct predictive scenarios that resemble real-world applications. For this purpose, we adopted the single-link clustering method, a bottom-up hierarchical clustering technique, ensuring that the sample distance between different clusters always exceeds a predefined distance, i.e., the minimum distance threshold *γ*. This guarantees high similarity within each cluster. To represent drug compounds, we utilized the binarized ECFP4 features, and for target proteins, we chose the integral PSC features. To accurately measure pairwise distances, we employed the Jaccard distance on ECFP4 and cosine distance on PSC. In the clustering of drugs and proteins, we set *γ*=0.5, a choice that avoids forming excessively large clusters and ensures the separation of dissimilar samples. The algorithm parameters we selected, including the single linkage clustering method, the binary ECFP4 features of drug compounds, the overall PSC features of target proteins, and the setting of *γ*, are all based on previous research [9, 24, 25], ensuring the accuracy and reliability of our method.

Regarding the model, the clustering-weighted module can be considered an optional component independent of the base learner. As depicted in Figure 1.e, we acquired cluster information of the training set through the aforementioned method and proportionally divided it into *k* datasets (where *k* being 5 in the figure). Subsequently, we selected *m* datasets (where *m* is 3 in the figure) to serve as the fine-tuning training set for the base learner. Concurrently, it’s necessary to construct a base learner and train it using the entire training set, and then fine-tune this base learner using the k derived fine-tuning datasets. When EADTN performs prediction tasks, we can look up the cluster information of the drug (target), and when fine-tuned base learners produce prediction scores, based on cluster affiliation, we enhance the credibility of the output from the relevant m fine-tuned base learners, resulting in the final prediction outcome. However, if we opt not to use this module, the model resorts to k-fold crossvalidation to establish *k* distinct training sets and train *k* different base learners. The final prediction is then derived using an average pooling method. The decision to use this module depends on the specific application context. To validate the performance of this module, we conducted a series of experiments across various application scenarios. The experimental results compellingly demonstrated the efficacy of this module and underscored the potential of ensemble learning in DTI prediction models.

## 3 Experiments

### 3.1 Evaluation strategies and metrics

Our experiments are based on two datasets: BindingDB [18] and BioSNAP [19]. For each experiment, the dataset is split into training, validation, and test sets at a ratio of 7:1:2, respectively. We evaluate the model’s performance in two experimental scenarios. Apart from the typical random splitting for predictions, we observed that in real-world applications, DTI models are often trained on data that is structurally or functionally similar to the drug (or target). To predict new interactions, and to emulate a genuine experimental prediction environment, we divided each cluster into training and test sets based on clustering information, and then merged the individual clusters. We refer to this scenario as ‘intra-cluster’ prediction.

To assess the performance of the model, we chose the Area Under the Receiver Operating Characteristic Curve (AUROC) and the Area Under the Precision-Recall Curve (AUPRC) as primary evaluation metrics, with accuracy, recall, and specificity as secondary metrics. Detailed explanations of these metrics are as follows:

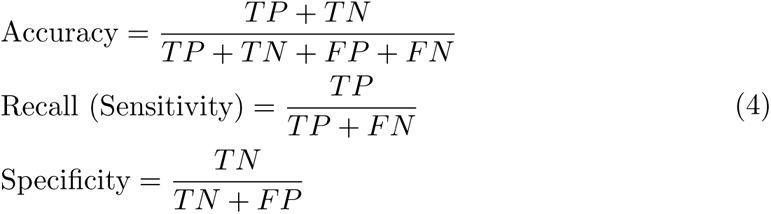

where TP and TN denote the number of positive and negative samples correctly predicted by the model; FP and FN denote the number of positive and negative samples incorrectly predicted by the model, respectively.

### 3.2 Experimental setting

In our experiments using the EADTN model, the following settings were adopted: batch size was set to 16; AdamW [26] was chosen as the optimizer; the learning rate was 1e-4; and weight decay was set at 1e-4. The model was allowed to run for a maximum of 200 epochs. To select the best-performing model, we made our choice based on the best Accuracy score on the validation set, using this score to assess the final performance on the test set. To prevent overfitting, we implemented early stopping with a patience value of 30. Regarding the feature encoder, both drug and target features were composed of three 1D CNN layers, with kernel sizes set to [4,6,8] and [4,8,12] respectively. The size of the embedding layer was 64. Additionally, we employed PolyLoss [27] as the loss function. The maximum sequence length was limited to 1,000 for proteins and 100 for drug molecules.

### 3.3 Model performance in random scenarios

We compared EADTN with six other methods. These include two machine learning algorithms: SVM [28] and RF [29], and four advanced deep learning methods: DeepConv-DTI [12], GraphDTA [11], MolTrans [10], and DrugBan [9]. Given that our experiment was set in a random scenario, we did not activate the clustering weighting module, and only applied average pooling to the outputs of the base learners.

Table 1 and Figure 3 show the performance of EADTN in comparison with six other baseline methods. Across the BindingDB and BioSNAP datasets, EADTN outperforms all the baseline models in every metric. It’s evident that increasing the number of base learners can further enhance the performance of the model.

**Fig. 3:**
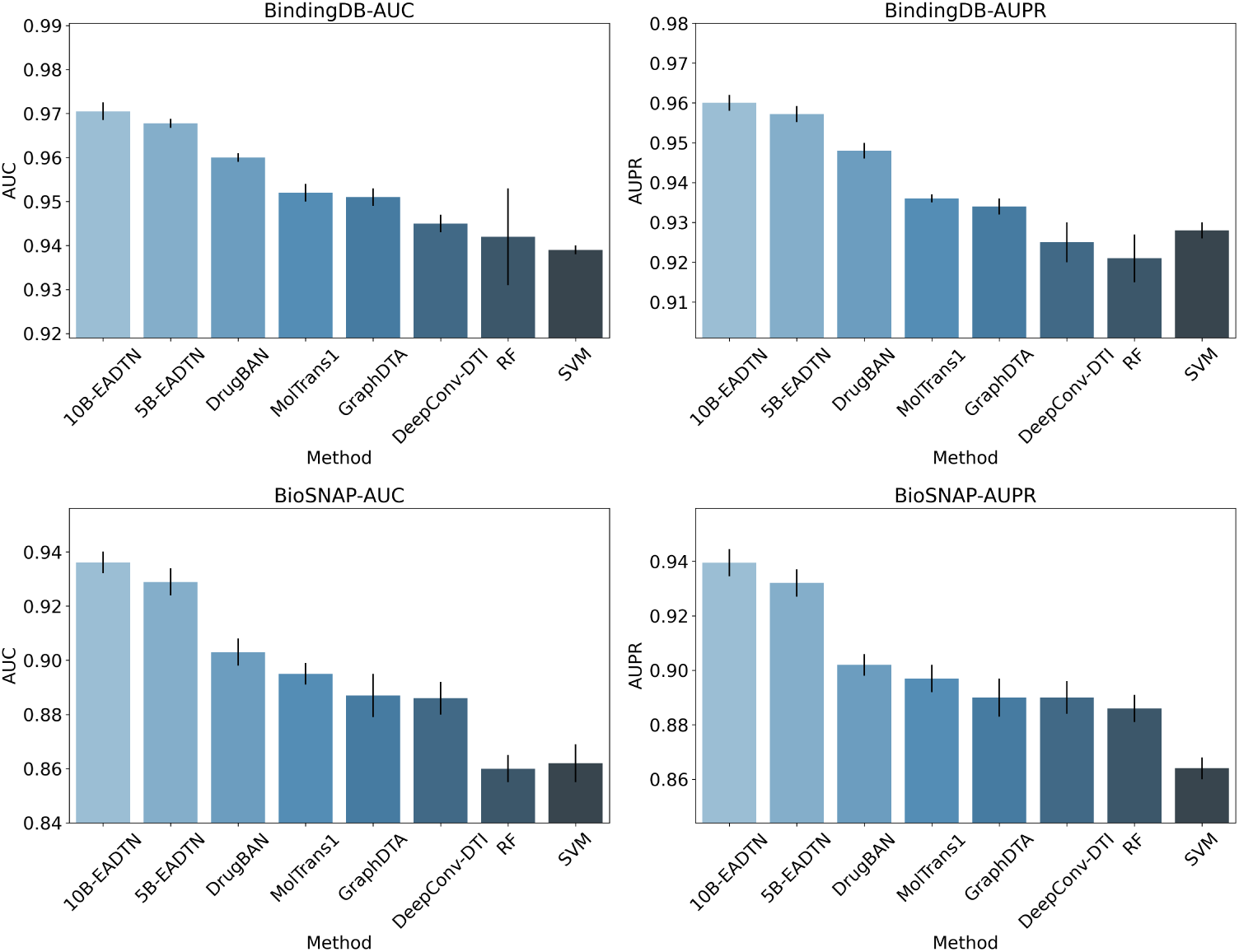
Comparison of AUC and AUPR Values Across Methods on the BindingDB and BioSNAP Datasets. Error bars in the chart represent standard deviations from the mean.

**Table 1:**
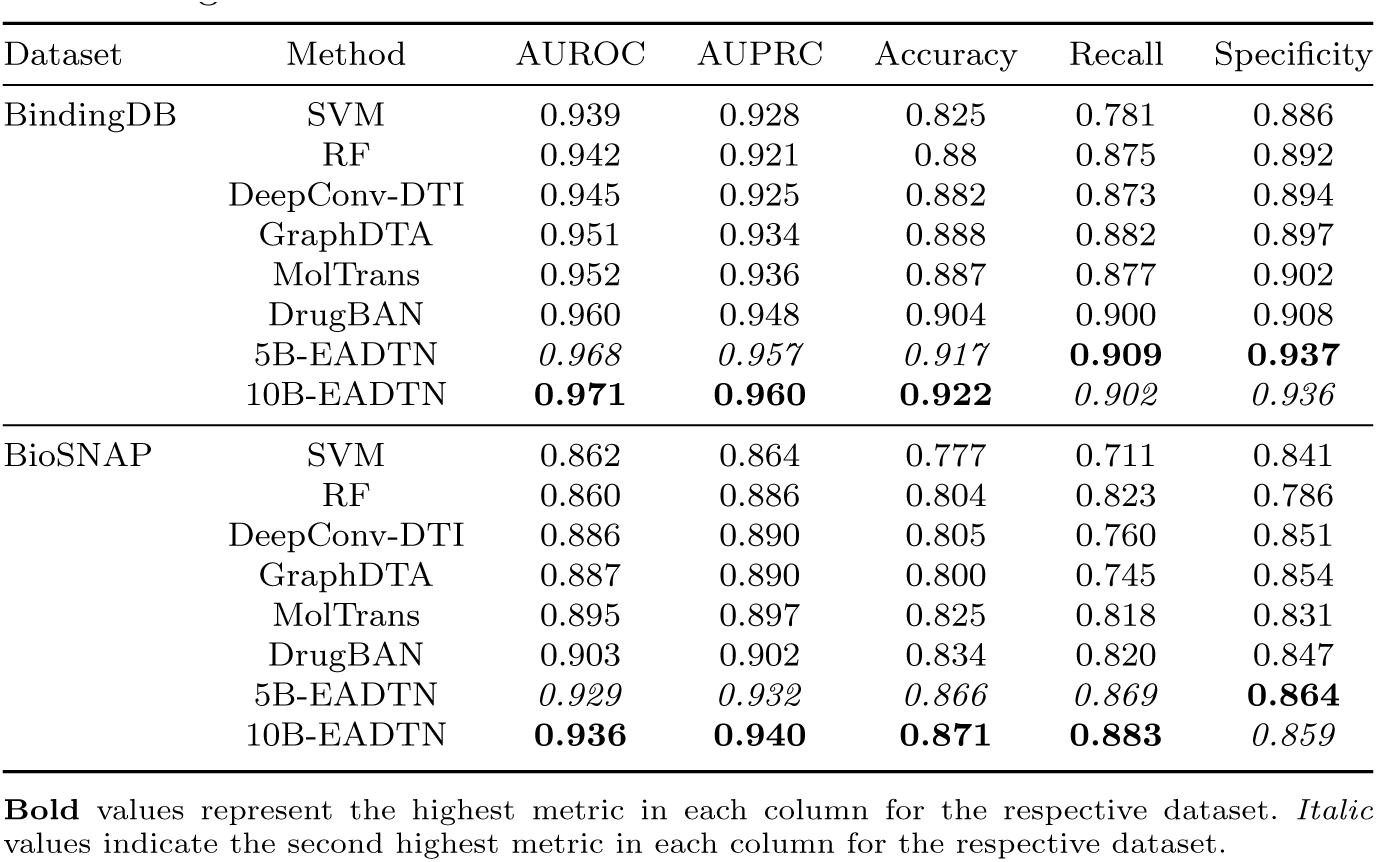
Comprehensive Performance Comparison of Various Methods on the BindingDB and BioSNAP Datasets.

**Table 2:** Performance Comparison of Various Methods in Intra-Cluster Scenarios with Drugs as the Clustering Criterion.

| Dataset | Method | AUROC | AUPRC | Accuracy | Recall | Specificity |
| --- | --- | --- | --- | --- | --- | --- |
| BindingDB | DrugBan | 0.96516 | 0.95154 | 0.90964 | *0.89163* | 0.93732 |
|  | 5B-EADTN | *0.96571* | *0.95535* | *0.92658* | 0.89014 | **0.95029** |
|  | 5B-EADTN-Fine-Tuning | **0.97471** | **0.96424** | **0.92908** | **0.90739** | *0.94318* |
| BioSNAP | DrugBan | 0.91419 | 0.92696 | 0.84567 | 0.83333 | 0.85682 |
|  | 5B-EADTN | *0.93375* | *0.94237* | *0.87144* | *0.86781* | **0.87845** |
|  | 5B-EADTN-Fine-Tuning | **0.94376** | **0.95298** | **0.87737** | **0.8823** | *0.87183* |

**Bold** values represent the highest metric in each column for the respective dataset. *Italic* values indicate the second highest metric in each column for the respective dataset.

**Table 3:** Performance Comparison of Various Methods in Intra-Cluster Scenarios with Targets as the Clustering Criterion.

| Dataset | Method | AUROC | AUPRC | Accuracy | Recall | Specificity |
| --- | --- | --- | --- | --- | --- | --- |
| BindingDB | DrugBan | 0.96199 | 0.94901 | 0.90125 | 0.88963 | 0.91834 |
|  | 5B-EADTN | *0.96398* | *0.95232* | *0.92852* | *0.89014* | **0.94349** |
|  | 5B-EADTN-Fine-Tuning | **0.96887** | **0.95832** | **0.92951** | **0.90682** | *0.92142* |
| Biosnap | DrugBan | 0.90748 | 0.92524 | 0.83873 | 0.84015 | 0.83744 |
|  | 5B-EADTN | *0.93004* | *0.94275* | *0.86889* | *0.86799* | **0.86997** |
|  | 5B-EADTN-Fine-Tuning | **0.94131** | **0.95281** | **0.86889** | **0.90347** | *0.84777* |

**Bold** values represent the highest metric in each column for the respective dataset. *Italic* values indicate the second highest metric in each column for the respective dataset.

On the BindingDB dataset, the 10B-EADTN version achieved scores of 0.971, 0.960, and 0.922 in key metrics like AUROC, AUPRC, and accuracy, respectively. The improvement in accuracy and specificity is comparable to the performance gap between basic machine learning techniques and the previously best-performing model. These metrics’ advancements suggest that the model has significantly enhanced its ability to identify negative samples. However, the increase in the number of base learners doesn’t significantly impact specificity. Thus, we are inclined to believe that EADTN’s adaptive feature weight network can discern the binding patterns between molecules and target proteins, allowing for accurate identification of negative samples.

On the BioSNAP dataset, the performance boost of the 10B-EADTN version is quite significant. The rise in key metrics such as AUROC, AUPRC, and accuracy all exceed 3.5%, indicating that EADTN’s improvement over the baseline models is comprehensive and meaningful. Notably, the Recall metric improved by 6% compared to the previous best model, suggesting that EADTN can learn a broader range of drugtarget binding patterns, providing more meaningful guidance for wet lab experiments. Additionally, with an increase in the number of base learners, the improvement in recall is also very noticeable. We believe this is because the DTI linkage relationships in this dataset are very dense. The multiple binding modes between drugs and targets make it difficult for other models to predict DTI relations across these varied patterns. However, EADTN, by creating diverse training data for ensemble learning, allows different base learners to capture these different DTI binding modes more effectively, significantly enhancing the model’s generalization capability.

EADTN’s performance in random scenarios suggests that its adaptive feature weight network can indeed capture more critical information and binding patterns between molecules and target proteins. The diversity offered by ensemble learning means the model can learn from a wider array of binding patterns. This further underscores the potential of ensemble learning in DTI prediction tasks and deep learning in general.

### 3.4 Model performance in intra-cluster scenarios

In drug repurposing [30] and the exploration and development of drug functions, interaction data of the drugs to be predicted hold decisive value for predictive models. Even though we might lack data for specific target drugs, drugs with similar structures can still provide ample interaction information for the model. However, most previous studies haven’t fully leveraged these structurally similar drugs and their relationships with targets to guide model predictions. Thus, we introduced an intra-cluster scenario. By clustering structurally similar drugs and targets, and then fine-tuning the base learners with the clustered training dataset, it allows for a deeper understanding of the binding patterns between similar drugs and targets. When the fine-tuned base learner provides prediction scores, based on cluster membership, we elevate the credibility of the corresponding fine-tuned base learner’s output, leading to the final prediction results. The outcomes demonstrate that, in this scenario, this method significantly enhances the predictive accuracy of the model.

On the BindingDB and BioSNAP datasets, 5B-EADTN-Fine-Tuning showed the best performance on all key evaluation measures. These results clearly show that our ensemble learning and fine-tuning approach greatly improves the model’s performance. Also, adding more base learners can give each one a more detailed area of focus. This means that each learner can better learn how similar compounds bind with their targets, making good use of information from similar drugs. This helps to make the model more reliable and accurate in its predictions. In future studies, we hope to use this method for more complex drug predictions to give even more accurate results.

Moreover, we delved deeply into the effects of the weight adjustments and the number of fine-tuning iterations on model performance. Specifically, we examined how the output weights of non-subordinate cluster-based learners contrasted with those of subordinate cluster-based learners in terms of influencing performance metrics. Using a baseline ratio of 1:1, we adjusted to 1:2 with an increment of 0.2. During this process, we closely observed the effects of 6 rounds of fine-tuning, with relevant results illustrated in Figure 4 and Figure 5.

**Fig. 4:**
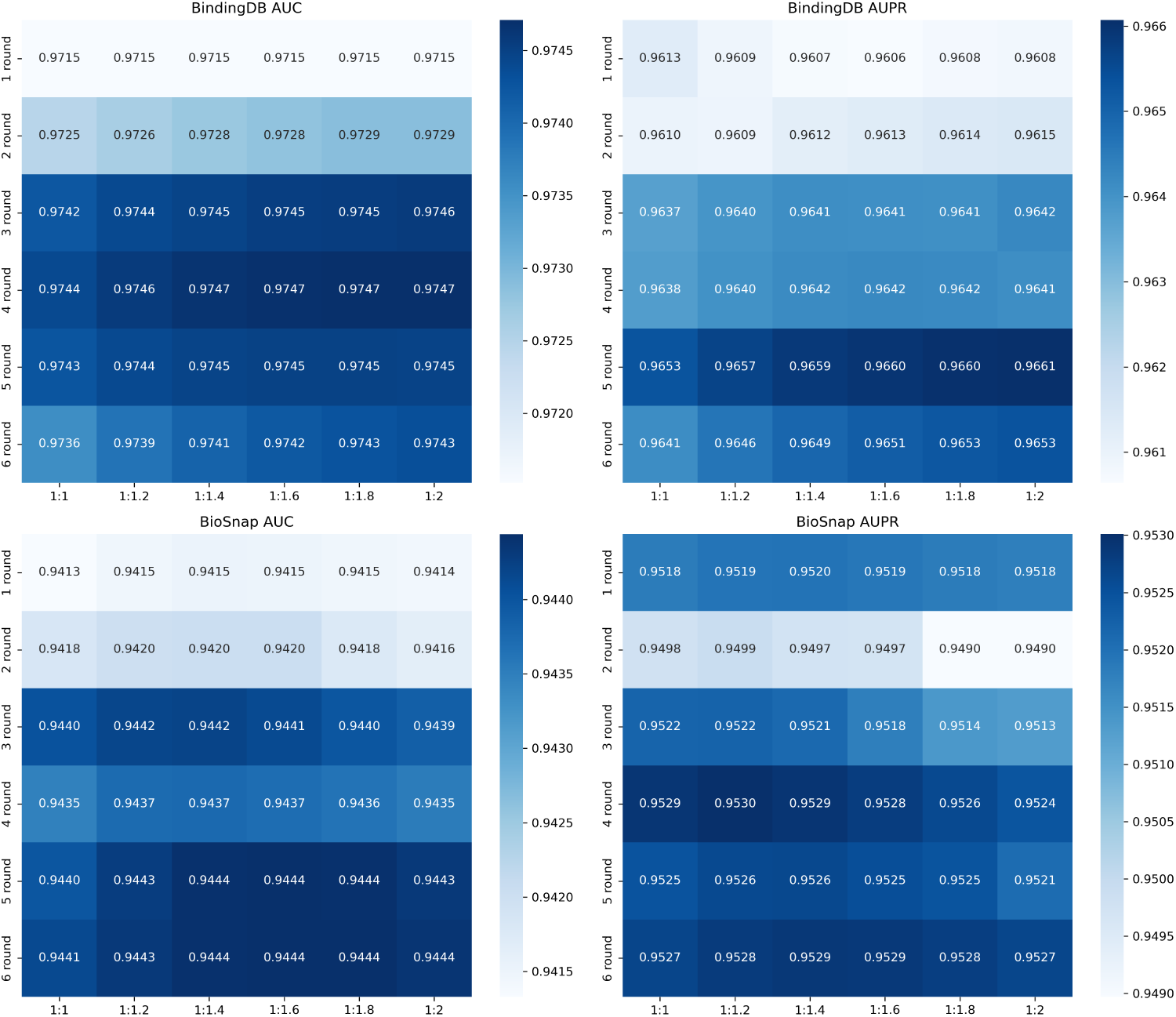
The Impact of Fine-Tuning Rounds and Weight Ratios on the Model’s Performance in Intra-Cluster Scenarios with Drugs as the Clustering Criterion.

**Fig. 5:**
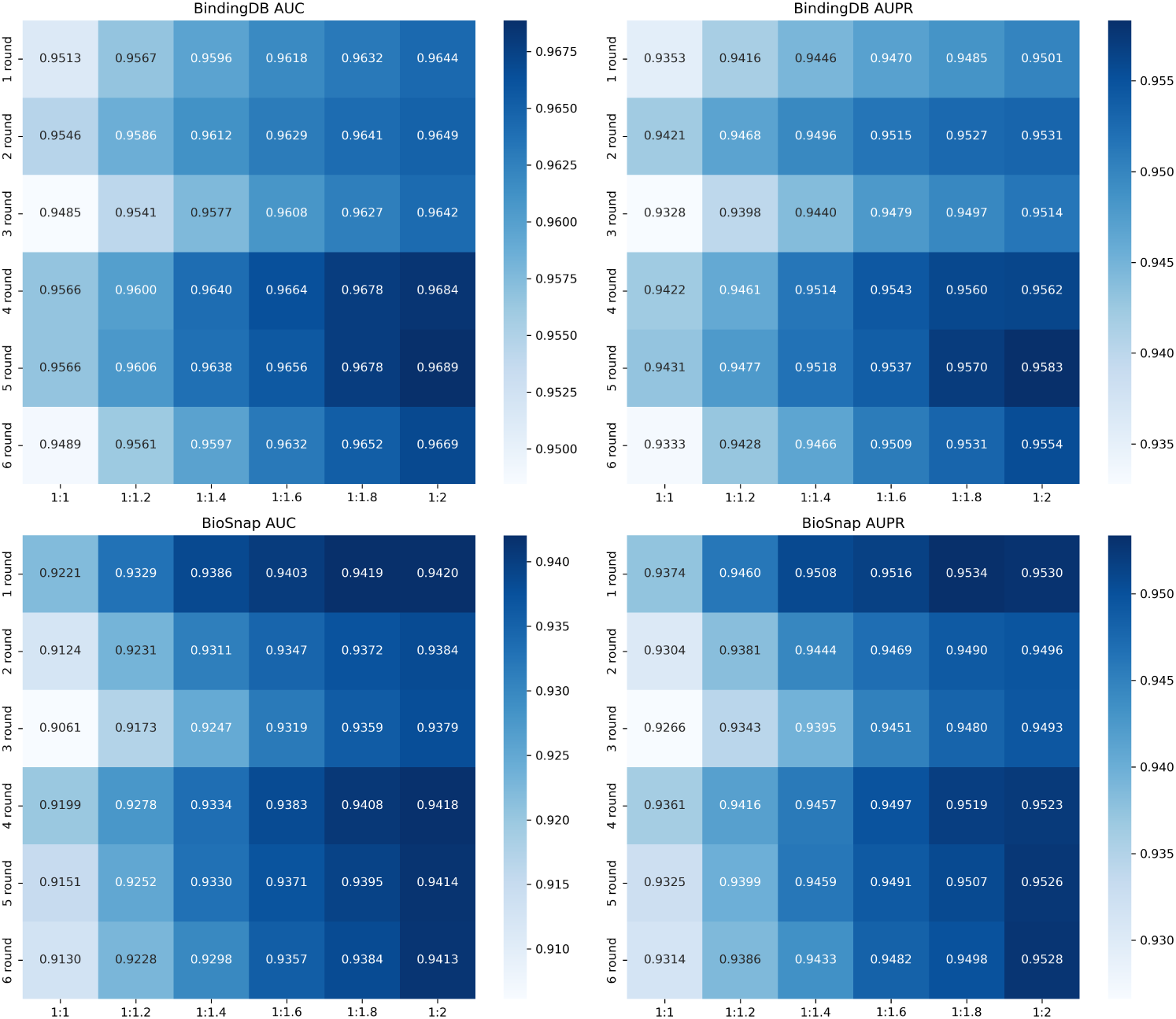
The Impact of Fine-Tuning Rounds and Weight Ratios on the Model’s Performance in Intra-Cluster Scenarios with Targets as the Clustering Criterion.

Based on the experimental findings, it is discernible that optimal performance is often achieved after 4-5 rounds of fine-tuning across both datasets. This is because excessive fine-tuning rounds might gradually erode the foundational knowledge acquired during the pre-training phase, thereby compromising the model’s credibility when predicting non-subordinate cluster DTI relationships. Concerning the weight ratio, we observed that in most cases, it is positively correlated with model performance. This observation further corroborates the enhanced reliability of finetuned base learners when predicting DTI relationships within the fine-tuned domain. However, the two clustering criteria have different impacts on the model. Schemes based on drug criteria are more sensitive to the number of training rounds, while those based on target criteria are more sensitive to the weight ratio. This could be attributed to the fact that, compared to protein targets, drugs from different clusters still exhibit certain similarities. As a result, even the outputs from base learners of non-subordinate clusters hold significant value. On the other hand, the differences between protein targets from different clusters are more pronounced, making the base learners more focused on the binding patterns of the subordinate clusters, and thus, the outputs from other base learners are less valuable. Moreover, it is worth noting that overly high weights could undermine the diversity of ensemble learning, leading to an oversight of the outputs from non-subordinate cluster learners, which in turn might diminish the model’s generalization capability.

## 4 Verification of the model’s ability to identify key substructures

In pursuit of a deeper understanding as to whether the EADTN model truly comprehends the interaction patterns between compound substructures and amino acid residues, we devised a methodology anchored on Shapley values, aiming to accurately gauge and rank the significance of each substructure. Shapley values, originating from game theory, were first introduced by Lloyd Shapley in 1953 and primarily delineate the contribution of each player in a cooperative game towards the total output [31]. In recent times, Shapley values have been expansively adopted in the realm of machine learning [32], especially in elucidating model decisions. Within the ambit of substructure importance analysis, conventional methodologies often perceive substructures as autonomous entities, determining their relevance based on model predictions. Such an approach tends to overlook the holistic structure of the compound and how individual substructures collaboratively influence molecular properties. This oversight becomes particularly pronounced in the context of drug-target interactions.

Taking into account the limitations of conventional techniques, our research adopts a fresh perspective: treating each substructure as players in a cooperative game, collectively determining the interactions between the molecule and the target. Herein, Shapley values are utilized to assess the marginal contribution of every substructure. We assign a “value” to each substructure, which represents the model’s prediction score for the interaction strength between the molecular structure and the target. Specifically, the SMILES of each substructure, when paired with a designated target protein, are input into the model. The resultant outputs from the decoder are then processed, subtracting the prediction value for ‘no interaction’ from that of ‘interaction’, yielding the “value” for the substructure. Subsequently, these values are employed to compute Shapley values, which in turn provide a ranking for the importance of each substructure. The comprehensive procedure can be referred to in Figure 6. Furthermore, to factor in the influence of the entire molecular structure on prediction outcomes, we retained the molecule’s backbone during substructure division. All substructures coexist based on this backbone. While the backbone imparts some foundational value to the substructures, its consistent presence in each marginal contribution calculation ensures it doesn’t skew our assessment of the marginal effects that substructures bring in comparison to the backbone. Thus, while keeping the backbone intact, we can dissect any part of the molecule to study the value different substructures bring to the specific properties of the molecule. The detailed computation protocol has been made publicly accessible in the code repository, with the calculation formula defined as follows:

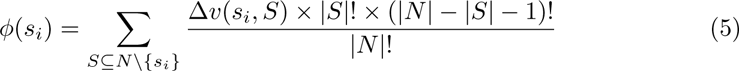

- *ϕ*(*s_i_*) represents the Shapley value of substructure *s_i_*.
- *S* is a subset of structures excluding *s_i_*.
- *N* is the set of all substructures.
- Δ*v*(*s_i_, S*) is the marginal contribution of substructure *s_i_* relative to set *S*, defined as Δ*v*(*s_i_, S*) = *v*(*S ∪ {s_i_}*) *− v*(*S*).
- *|S|* and *|N|* denote the sizes of sets *S* and *N* respectively.

**Fig. 6:**
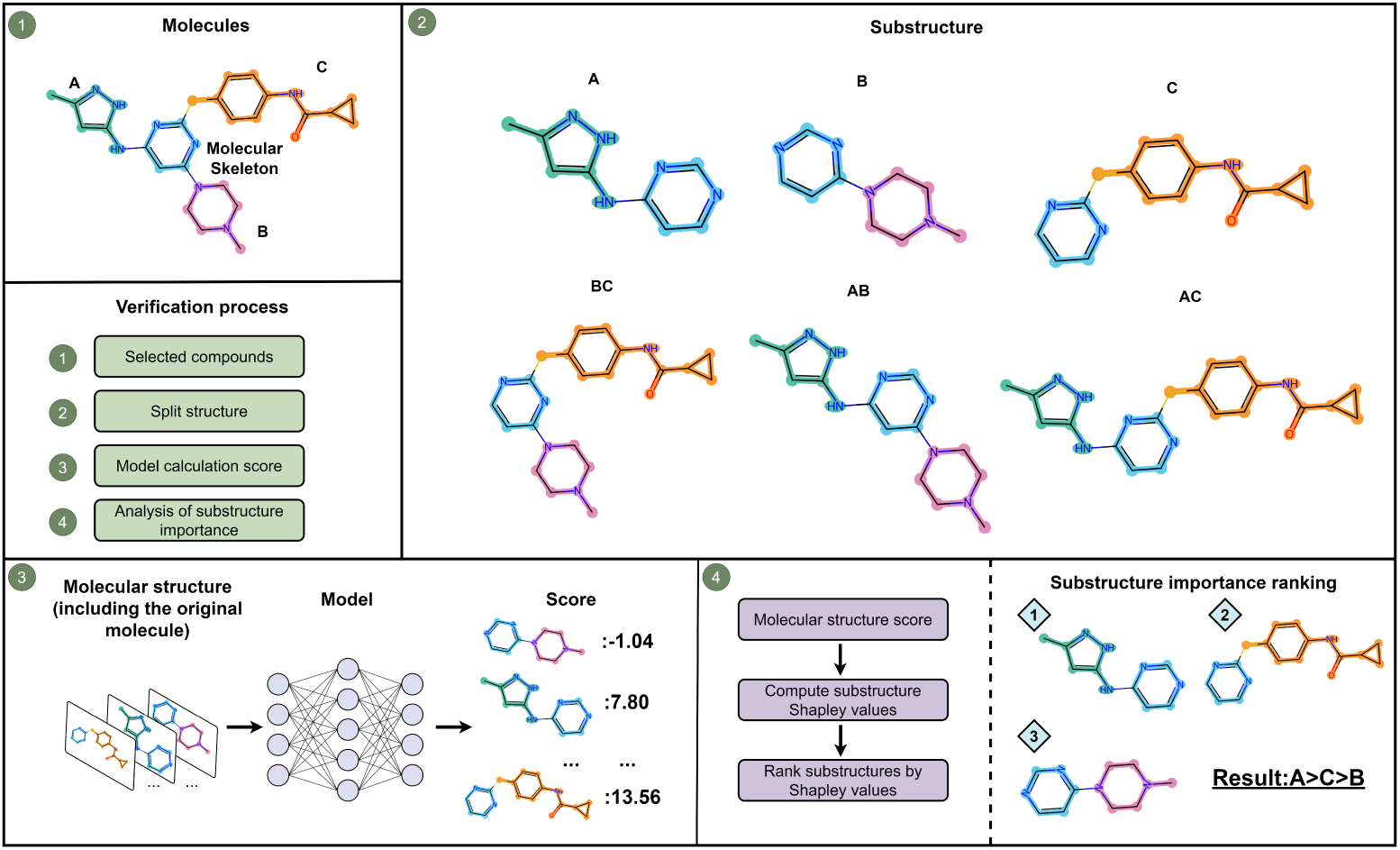
Algorithm Flowchart for Ranking Substructure Importance. The procedure for identifying and ranking substructure importance is as follows:

1. Select the target compound.
2. Form all possible sets by combining substructures, for the purpose of computing marginal contributions.
3. Feed the substructures into the model. Use the model’s prediction scores as the value for each set.
4. Calculate the Shapley value for each substructure based on the values of all sets, and subsequently rank them.

**Fig. 7:**
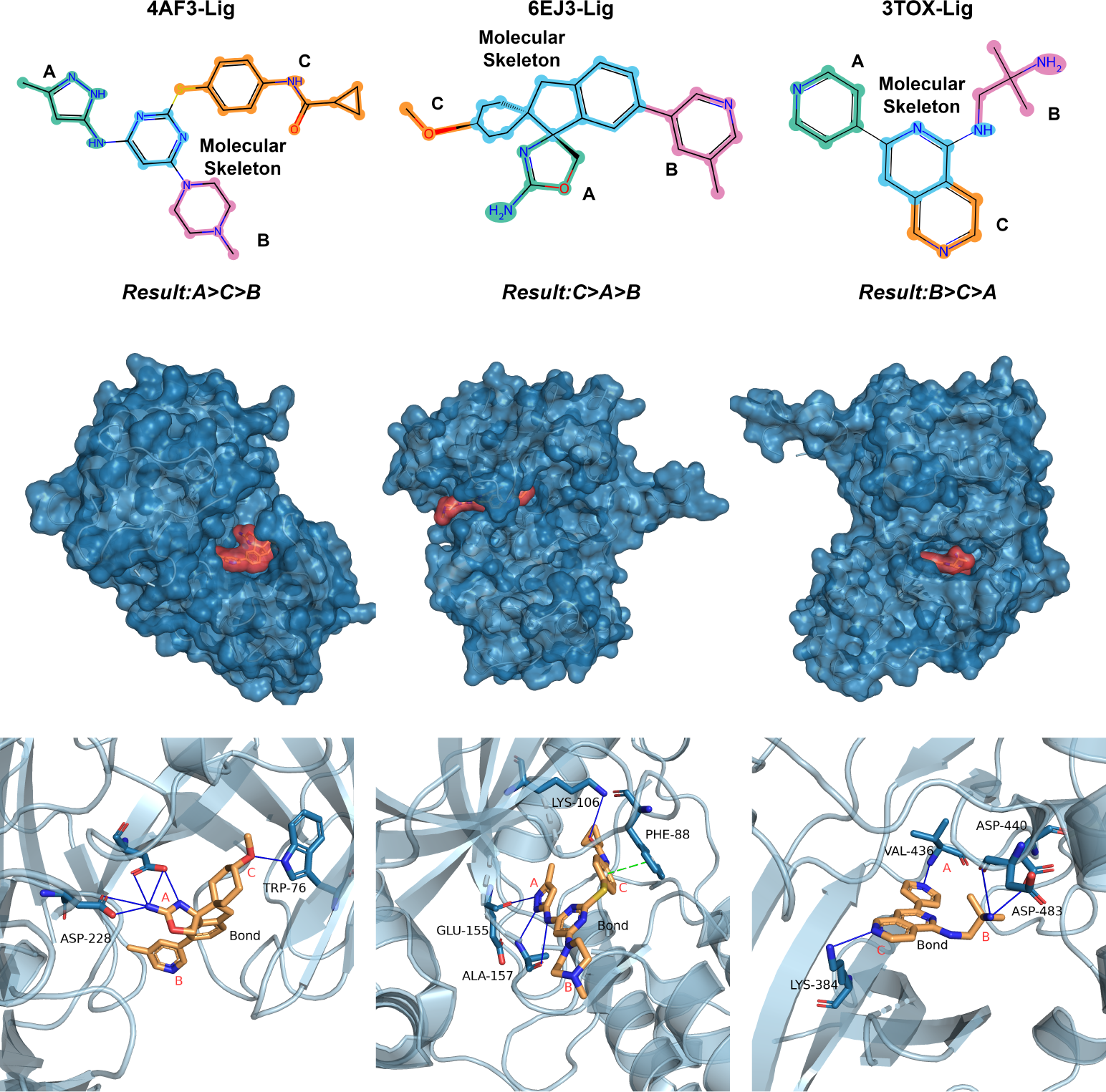
Precision Molecular Visualization of Substructure Verification. In the molecular surface representations of the second row, the target protein is colored in blue, and the drug is in red. In the images of the third row, amino acid residues involved in hydrogen bond and*π − π* stacking interactions with the drug molecule are explicitly labeled. Blue solid lines represent hydrogen bonds, while green dashed lines indicate *π − π* stacking interactions.

In order to assess the model’s capability in recognizing key substructures, we examined three targets from the BindingDB dataset (PDB ID: 4AF3 [33], 6EJ3 [34], 3TXO [35]). Corresponding ligand molecules, which had been experimentally validated and were not present in the training set, were sourced from RCSB [36]. For the evaluation of substructure significance, we primarily focused on hydrogen bonds and *π – π* stacking interactions as criteria and conducted inspections in the molecular operation environment [37], the specific visualization results are shown in Figure **??**.

For the PDB structure 4AF3 [33] (Human Aurora B Kinase in complex with INCENP and VX-680), our model adeptly identified the substructure 2-methylpyrazole (labeled as A) as the most significant. Experimental validation in the molecular environment revealed that substructure A forms three hydrogen bonds with the glutamic acid and alanine residues of the protein. Additionally, the substructure benzylpropanamide (labeled as C) engages in both a hydrogen bond and a *π – π* stacking interactions, and is correctly identified as the second most important sub-structure. In structure 6EJ3 [34] (BACE1 compound 23), the model perceived the ether link (labeled as C) to be of utmost significance. However, in reality, substructure C forms only a single hydrogen bond, whereas the imidazoline derivative (labeled as A) forms multiple. While the model did not precisely elucidate this fact, it successfully ranked 2-methylpyrazole as the least significant substructure. In the structure 3TXO [35] (PKC eta kinase in complex with a naphthyridine), the model adeptly identified the aminomethyl group (labeled as B) as the most pivotal substructure. Both substructures A and C form an equivalent number of hydrogen bonds. Further investigation into the weaker hydrophobic interactions revealed that substructure A possesses a marginally larger hydrophobic surface area compared to substructure C, and its hydrogen bond lengths are also shorter.

In conclusion, EADTN exhibits a commendable reliability in interpreting key sub-structures. Although it may not always precisely rank certain substructures, its ability to identify pivotal substructures is invaluable. This indicates that EADTN has successfully learned the interaction patterns between compound substructures and amino acid residues, further affirming its exemplary performance. Moreover, the methodology of ranking substructure significance based on shapley values is model-agnostic, paving the way for researchers to further evaluate various models in the future.

## 5 Case study

In the context of databases pertinent to drug-target interactions, positive samples are typically documented, while negative samples are predominantly generated through negative sampling. Such an approach may lead to an elevated rate of false negatives, omitting certain valuable interaction relationships. Although current strategies often employ data augmentation and semi-supervised learning to address this challenge, the efficacy of these methods remains limited. Nevertheless, EADTN, by leveraging cross-validation and ensemble learning, successfully mitigates the misleading effects introduced by false negative samples. To demonstrate the superiority of EADTN in this aspect, we sorted samples labeled as negative in the Biosnap dataset in descending order based on their scores, and selected the top four drug-target combinations. Furthermore, we utilized AutoDock [38] for molecular docking, verifying the genuine interaction circumstances of these negative samples from a molecular structure perspective based on docking binding energy values as well as the quantity of hydrogen bonds and *π−π* stacking interactions. The tabel illustrates the blind docking outcomes of the selected ligands and their respective targets. Comprehensive docking results can be accessed in our Github repository.

From an evaluation metric standpoint, a smaller binding energy (negative value) indicates a more potent interaction between two molecules. Many researchers opine that a binding energy less than −5.0 kcal/mol might suggest a strong interaction between the molecules in specific scenarios. Moreover, hydrogen bonds and *π – π* stacking interactions are two prevalent types of non-covalent interactions that play a pivotal role in the stability of molecular binding. While binding stability also encompasses other non-covalent interactions, binding pocket specificity, etc., the presence of two or more hydrogen bonds and at least one *π −π* stacking interaction often suggests a high likelihood of intermolecular interactions.

The experimental results are shown in Table 4 and Figure 8, all five drugs and their corresponding targets exhibited strong mutual interactions at the molecular structural level. Especially considering the binding energy criterion, the binding energies of all drug-target combinations were less than −5.0 kcal/mol, implying effective intermolecular interactions. Specifically, the binding energy of Bosutinib with 1WSR reached −7.5 kcal/mol, the lowest among all combinations. Coupled with its four hydrogen bonds and one *π − π* stacking interaction, it is inferred that this drug-target binding is particularly tight and stable. Other drug-target combinations, like Melatonin with 4HCT, Tamoxifen with 2HE3, and SU11652 with 2HQU, also exhibited binding energies less than −6.0 kcal/mol, accompanied by two or more hydrogen bonds, further corroborating their effective binding.

**Fig. 8:**
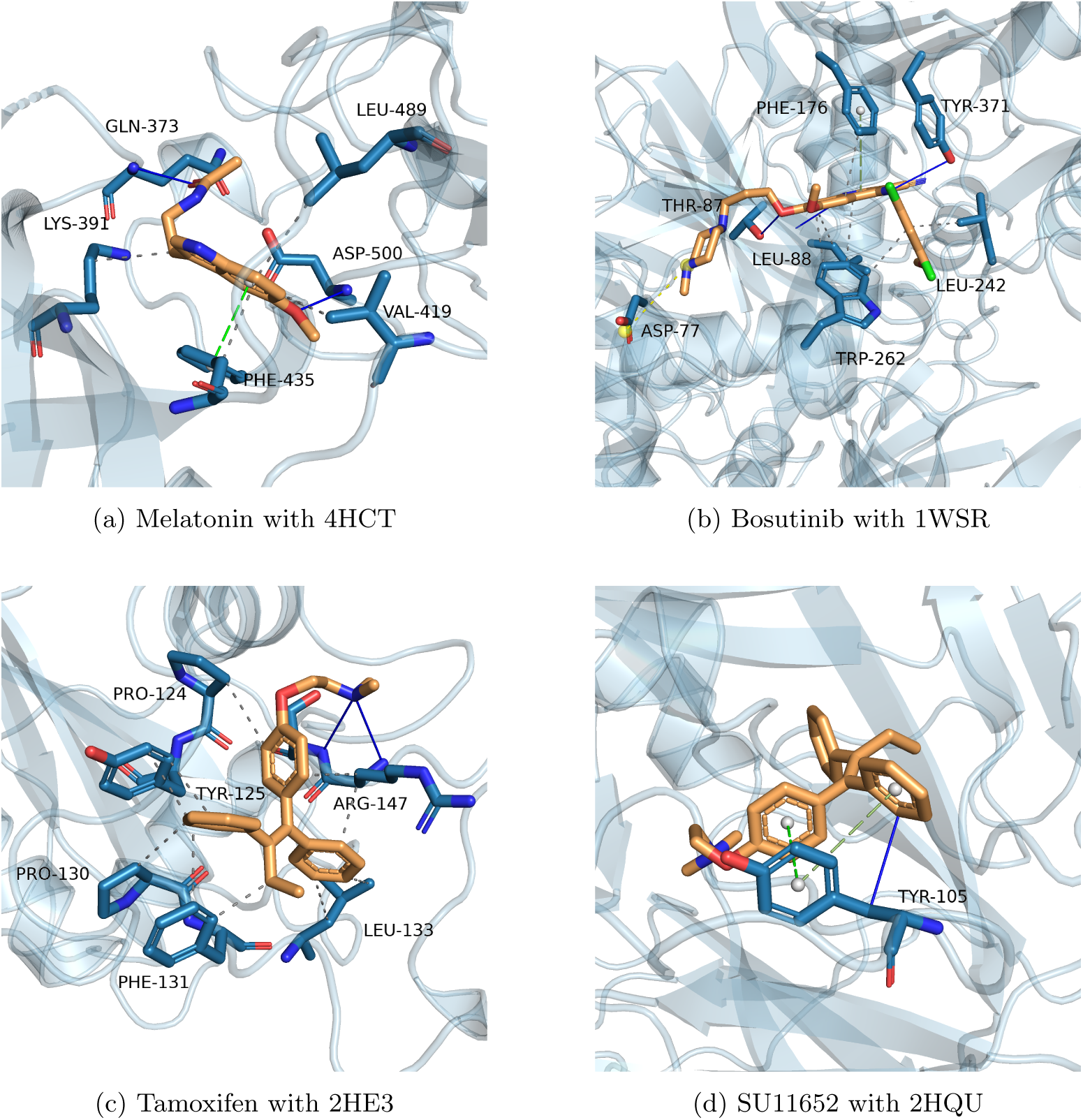
Molecular visualization of case study. The blue solid line represents hydrogen bond interactions, the green dashed line indicates *π−π* stacking interactions, the white dashed line signifies hydrophobic interactions, and the yellow dashed line denotes salt bridge (or ionic bond) interactions.

**Table 4:** Molecular docking verification of the top 4 negative samples.

| Rank | Drug | PDB | Energy (kcal/mol) | H-Bonds | $\pi - \pi$ stacking interactions |
| --- | --- | --- | --- | --- | --- |
| 1 | Melatonin | 4HCT [39] | -6.8 | 2 | 1 |
| 2 | Bosutinib | 1WSR [40] | -7.5 | 4 | 1 |
| 3 | Tamoxifen | 2HE3 | -6.5 | 2 | 0 |
| 4 | SU11652 | 2HQU [41] | -7.0 | 1 | 2 |

In summary, EADTN has demonstrated significant efficacy in reducing the misleading effects induced by false-negative samples. This case underscores potential shortcomings in traditional drug-target interaction databases and further attests to the unique value of EADTN in uncovering overlooked interactions. Such discoveries not only offer tangible insights for drug development and disease treatment but also instigate profound contemplation on optimizing the utilization of existing data resources.

## 6 Conclusion and discussion

In this work, we propose EADTN, a deep learning model based on adaptive feature weights and ensemble learning. We use the adaptive weight module to capture the significant relationship between protein residues and drug atoms, aiming to enhance the model’s generalization ability and interpretability. Concurrently, we employ an ensemble learning strategy to fully harness the diversity of the data, aiming to further reduce the model error. Moreover, we implemented a cluster fine-tuning strategy, which further enhances the model’s role in drug repositioning and new drug development scenarios. Compared to other state-of-the-art deep learning models, EADTN has demonstrated clear advantages across various evaluation metrics.

To explore the model’s potential in substructure recognition, we designed a substructure importance evaluation method based on Shapley values, further attesting to EADTN’s exceptional performance in substructure identification tasks. Additionally, although negative sampling of the data still poses challenges due to false negatives, we effectively mitigated this issue through cross-validation and ensemble learning strategies. Case studies further substantiate our claims.

Overall, the focus of this study is on leveraging the diversity of ensemble learning, using multiple base learners to reduce generalization error. By building upon these base learners for fine-tuning, the overall performance of the model is elevated. This approach paves new avenues for the application of ensemble learning in related domains. We believe that by continuing to expand our research in this direction, it can offer novel insights for other fields.

## 7 Data and Code Availability

Our computational models and corresponding code are accessible via: https://github.com/kkkayle/EADTN.

